# Using DNA origami to study nanoscale organization of plasma membranes

**DOI:** 10.1101/2025.08.27.672545

**Authors:** Eloina Corradi, Konlin Shen, Zeynep Karatas, Maureen Cercy, Thomas Schlichthaerle, Margaux Caumont, Mélissande Osouf, Brune Vialet, Philippe Barthelemy, Morgane Rosendale, Adiyodi Veetil Radhakrishnan, Tianchi Chen, Ralf Jungmann, Arnaud Gissot, Shawn M. Douglas, Grégory Giannone

## Abstract

Plasma membrane (PM) lipids and proteins are organized into nanoscale regions called nanodomains, which regulate essential cellular processes by controlling local membrane organization. Despite advances in super-resolution microscopy and single particle tracking, the small size and temporal instability of nanodomains make them difficult to study in living cells. To overcome these challenges, we built fluorescent DNA origami probes that insert into the PM via lipid anchors displayed on the cell. The number and spatial distribution of anchors between the origami and the cell surface were precisely defined by the origami, enabling nanometer-scale sampling of the cell surface. Inserting these DNA origami particles into the membrane with lipid anchors allowed them to passively diffuse across the membrane, and we tracked their movement using single particle tracking to survey the PM landscape. By varying the number and spatial arrangement of lipid anchors connecting the DNA origami to the cell surface, we showed that immobilization of DNA origami particles requires simultaneous interactions with multiple nanodomains. Disruption of the actin cytoskeleton reduced immobilization, confirming its role in supporting nanodomain stability. Moreover, transient mechanical stretching of cells led to reversible increases in DNA origami mobility, indicating that mechanical force can reversibly regulate PM nanodomain organization. Altogether, we present a novel membrane-integrated DNA origami approach that provides mechanistic insights into PM nanodomain architecture and dynamics in living cells.

## Introduction

The plasma membrane (PM) separates the cell interior from the extracellular environment and is a crucial hub for numerous cellular processes including signal transduction, intercellular communication, integration of mechanical signals, control of cell adhesions, and cytoskeleton organization ^1,2^. The functional role of the PM largely relies on the lateral heterogeneity of its components (i.e. lipids and proteins), which are locally concentrated to generate functional domains of different sizes and lifespans ^3^. Although some domains can span micrometers, they are fundamentally composed of nanoscale structures (10–200 nm), called nanodomains, that compartmentalize and coordinate membrane-based activities ^4,5^. PM nanodomains regulate a wide range of functions through multiple mechanisms. For instance, protein clustering in PM nanodomains regulates key signaling pathways involving Ras small GTPases ^6^, Src-family kinases ^7^, and interferon-γ receptor (IFN-γR)^8^. PM nanodomains also contribute to several cellular processes including immunogenetic response to cancer ^9^, cell adhesions ^10^, cell migration, ^11^ and phagocytosis ^12^ through transmembrane proteins such as integrins ^10,11^ and CD44 ^12,13^ that link the extracellular matrix (ECM) to the actin cytoskeleton. PM nanodomains also control PM topography, enabling cargo endocytosis ^14^ and viral budding ^15^, and are involved in mechanotransduction ^11,16^. The distribution, composition, and dynamics of PM nanodomains enable precise spatial and temporal control of various cellular functions yet we know little about how they accomplish these complex tasks.

Our understanding of how PM nanodomains coordinate cellular processes has been limited by the challenges of visualizing these small, dynamic structures. Current insights into PM nanodomain organization primarily rely on advanced optical techniques such as super-resolution microscopy and single-particle tracking (SPT), as no other optical technique has sufficient spatiotemporal resolution to accurately characterize PM nanodomains ^5^. To study nanodomain organization and dynamics, an ideal technique would combine high spatial and temporal resolution with the ability to sample large areas of the plasma membrane in living cells. However, current methods typically meet only some of these requirements. Super-resolution imaging of fixed samples offers high spatial resolution and wide sampling areas, but cannot capture dynamic behavior ^16,17^. Near-field single-molecule optical microscopy (NSOM) ^10^ and homo-FRET ^18^ achieve nanometer-scale precision (∼5 nm), either directly through near-field optical scanning or indirectly via energy transfer efficiency, but are limited by low temporal resolution (seconds) and, for NSOM, by small fields of view. In contrast, single-particle tracking methods significantly improve temporal resolution (down to milliseconds) while maintaining high localization precision (2-30 nm). Yet, this comes at the cost of reduced sampling area (e.g. STED-FCS ^19,20^) and limited molecular coverage, as they typically track individual proteins (e.g. GPI-anchored proteins) ^7,21,22^ or lipids (cholesterol, sphingolipids) ^19,20,23–25^, providing only partial insight into overall nanodomain organization.

The DNA origami method folds a long single-stranded DNA (ssDNA) scaffold into a user-defined structure with single-base precision, resulting in a nanoparticle that can be sculpted with single-nanometer resolution ^26^. They have been used to unveil and control ligand–receptor mechanisms by spatially organizing either receptors ^27,28^ or ligands ^29–31^ with nanometric precision. Furthermore, DNA nanostructures have been patterned with lipophilic anchors to interface with and shape both model ^32–36^ and cellular membranes ^37,38^, for instance to induce deformation ^33,36^, curvature ^34^, perforation ^36,38^ or protection ^37^. Despite their promise as molecular probes, DNA origami have not been applied to investigate plasma membrane nanodomain organization and dynamics in living cells. On this basis and building on our previous DNA origami pegboards ^27,28^, we developed an innovative approach based on DNA origami. This approach enables the study of nanodomain organization and dynamics in living cells with both high temporal resolution (millisecond range) and spatial localization precision (∼30 nm), while simultaneously sampling larger areas (38.5 nm × 36 nm) of the PM. We patterned and extended ssDNA handles off a DNA origami chassis to interrogate the organization of PM nanodomains through hybridization with complementary ssDNA-conjugated lipid anchors inserted into the PM. The precise spatial arrangement of our handles enables interrogation of membrane features smaller than 30 nm, far lower than the optical resolution limit. We decorated the origami chassis with multiple fluorophores to ensure a longer-lasting fluorescent signal without compromising the field of view. By embedding these DNA origami probes into cell PMs and measuring their diffusion characteristics using single particle tracking ^39–41^, we provide new insights into the nanoscale organization of PM nanodomains and their response to global perturbations such as actin destabilization and cell mechanical forces.

## Results

### DNA origami tracking can be used to probe cell membrane

Our PM nanodomain probes were based on a “molecular pegboard” design previously used to probe T cell activation ^28^ and phagocytosis ^27^. We folded an 8064-nt ssDNA scaffold into a roughly 60 nm x 60 nm x 6 nm rectangular prism. On one side of the structure, we added 12 biotin sites for conjugating monomeric-Streptavidin labelled with ATTO594 fluorophore (mStravATTO594). On the other side of the structure, we designed 72 uniquely-addressable sites where a 16-nt ssDNA handle for probing the membrane could be extended. The sites were separated from each other by 3.5 nm in one direction and 7.2 nm in the other direction, allowing for PM interrogation at the nanometer scale (Fig.1a, Fig.1b).

**Figure 1.**
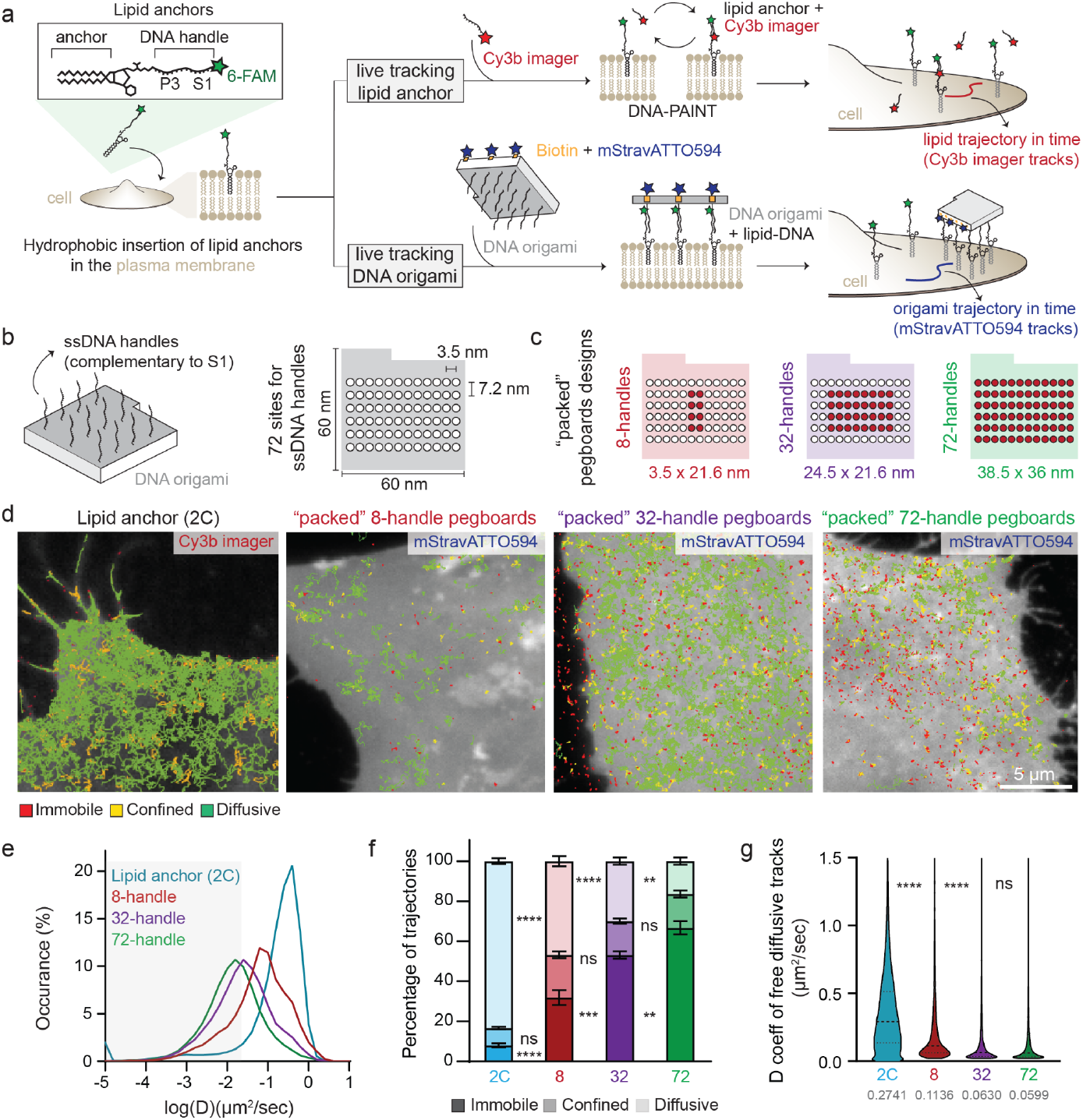
DNA origami diffusion in the PM depends on the number of handles for lipid anchors. **a)** Schematic of the experimental strategy: DNA origami displaying arrangements of ssDNA handles are hybridized to synthetic lipids anchors displayed on cell membranes. Using single particle tracking, we measure the diffusion characteristics of the DNA origami across the cell surface to probe the nanoscale landscape of the plasma membrane in living cells. Single lipids are tracked through hybridization with a P3 Cy3b imager. **b**,**c)** Schematic of the DNA origami pegboard (b) and of the three “packed” design with 8, 32 or 72 ssDNA handles for lipid anchors (c). **d)** Trajectories of lipid anchors or DNA origami overlaid on 6-FAM cell plasma membrane (gray). Trajectories are color coded to show their diffusion modes: diffusive (green), confined (yellow) and immobile (red). **e)** Distributions of the mean diffusion coefficient D computed from the trajectories of lipid anchors 2C or DNA origami. **f)** Fractions of tracked lipid anchors 2C and DNA origami undergoing free diffusion, confined diffusion, or immobilization in the plasma membrane. **g)** Diffusion coefficient D for all free diffusive tracks of the lipid anchors 2C and DNA origami. Median values reported. Data information: mean ± SEM (f), median and interquartile range (g). Statistics: n=5 (2C), n=6 (8-handles), n=7 (32-handles), n=6 (72-handles), e) 2-way ANOVA with Tukey multiple comparisons test, f) number of diffusive tracks 132543 (2C), 7312 (8-handles), 8487 (32-handles), 14864 (72-handles), Kruskal-Wallis test with Dunn’s multiple comparisons test. Abbreviations: 2C, lipid anchor with two aliphatic chains.

To enable interactions between the cells and the pegboards, we decorated the cells with lipid anchors conjugated to 34-nt ssDNA strands (Fig.1a). The first 16 bases of the lipid-anchor ssDNA strands are complementary to the ssDNA handles displayed on the DNA origami pegboard (S1 site, Fig.1a, Fig.S1a). The length and sequence of the handles were chosen to minimize secondary structure formation as well as to stabilize duplex formation at room temperature. The last 11 bases on the 3’ end of the lipid-anchor strand can hybridize to an “imager” strand, carrying a Cy3b fluorophore (P3 site, Fig.1a, Fig.S1a). The 11-base sequence was designed for transient binding to allow live tracking of the lipid anchors via DNA-PAINT (Fig.1a).

Previous work with DNA origami nanostructures on lipid membranes have shown that the diffusion characteristics are dependent on the lipid composition ^42^. To determine the type of lipid anchor to use, we synthesized two different lipid anchors bearing long saturated aliphatic chains, components characteristic of PM nanodomains ^43^: one with a single 18-carbon tail (1C), and another with two 15-carbon tails (2C) (Fig.S1a,b).

Lipid anchors were conjugated to 6-carboxyfluorescein (6-FAM) at the 3’ end of ssDNA to monitor PM incorporation in cultured mouse embryonic fibroblasts (MEFs) (Fig.S1a). Fluorescence imaging showed that both 1C and 2C lipid anchors were incorporated into the outer leaflet through hydrophobic insertion (Fig.S1c). Addition of Cy3 imager strands enabled tracking of thousands of trajectories for both lipid anchors (Fig.S1c). Control experiments without lipid anchors showed no detectable tracks, confirming trajectory specificity. For trajectories exceeding 200 ms (>10 points), we computed the mean squared displacement (MSD) and diffusion coefficients (D) (Fig.S1d). By fitting the MSD over time, we classified diffusion modes of lipid anchors as immobile, confined, or free diffusive ^38^ (Fig.S1e). MSD analysis revealed that both types of lipid anchors predominantly diffuse freely within the membrane, with minimal confined or immobile fractions (Fig.S1c-f). The 2C lipid anchor exhibited higher diffusion coefficients than 1C anchor, indicating faster free diffusion (Fig.S1f). Validation experiments at 500 Hz acquisition confirmed free diffusion dominance and the faster 2C versus 1C dynamics (Fig.S1g-j). These results establish that individual synthetic lipids behave similarly to native saturated lipids ^19,20,25^, providing a foundation for DNA origami-based nanodomain organization studies.

We next confirmed that DNA origami pegboards diffuse on the PM when bound to lipid anchors. Using a test DNA origami structure that had 12 DNA handles, we tracked pegboards during live imaging. Without lipid-anchor insertion in the PM, no origami tracks were detectable on cells. However, once we decorated the cells with lipid anchors, the DNA origami attached to the cell membranes and could be tracked (Fig.S2a). Compared to individual lipids, the DNA origami particles exhibited decreased free diffusion and increased immobilization, regardless of having 1C or 2C lipid anchors (Fig.S2b-d). Unlike individual lipids, the DNA origami did not exhibit significantly different diffusion coefficients or fraction of diffusion modes when bound to 1C or 2C lipid anchors (Fig.S2c,d), likely due to dominant viscous drag effects in the membrane. We selected 2C lipid anchors for subsequent experiments, matching the dual aliphatic chain structure typical of PM lipids. These results demonstrate that, while individual lipids remain largely free-diffusing, DNA origami bound to multiple lipid anchors display confined diffusion and immobilization on the PM, thus confirming the feasibility of the proposed strategy to probe nanoscale organization of the PM.

### Pegboard diffusion in the PM depends on the number of handles for lipid anchors

The increased immobilization and slower diffusion of the DNA origami pegboards compared to single lipids could be due to genuine interactions with lipid anchors within PM nanodomains, interactions with native extracellular components of the PM, unintended aggregation of the pegboards, or some combination of all three. To differentiate possible effects of these mechanisms, we varied the number of handles per pegboard (Fig.1c). If diffusion behavior remained unchanged, pegboard-pegboard or pegboard-extracellular interactions were likely providing the dominant effect on immobilization. Conversely, if diffusion behavior correlated with handle number, this finding would indicate that larger membrane engagement areas increase the probability of engaging with PM nanodomains, enabling detailed spatiotemporal exploration.

We generated three pegboard variants with either 8, 32, or 72 handles, designed to bind lipid anchors across 3.5 nm × 21.6 nm, 24.5 nm × 21.6 nm and 38.5 nm × 36 nm areas, respectively (Fig.1c). After incorporating 2C lipid anchors into MEF cell membranes, we added pegboards and quantified their diffusive behavior using single particle tracking (SPT) (Fig.1d). Higher handle numbers correlated with increased immobilization: 8-handle pegboards showed the highest diffusive trajectory percentage, followed by 32-handle and 72-handle pegboards (Fig.1d-f). Diffusion coefficients decreased with handle number (Fig.1e,g). These results demonstrate that pegboard diffusion on the PM depends on the number of lipid anchor interactions with membrane components rather than on the DNA origami pegboard itself, indicating that observed dynamics reflect membrane properties. This validates our DNA origami strategy as a promising tool to better understand the organization of PM nanodomains.

### Spatial arrangement of lipid anchors influences DNA origami diffusion

We wondered if the observed changes in the pegboard’s diffusive properties stem solely from increased membrane drag triggered by a higher number of handle-lipid anchors. To determine whether we were probing PM organization or simply seeing the effect of more points of contact between the origami and the cell PM, we limited the pegboard handle number to 8, but varied the spatial arrangement of the handles to change the membrane footprint of the pegboards, alongside a 72-handle control (Fig.2a,b). After inserting lipid anchors into cell membranes and mixing with pegboards, we tracked pegboard diffusion along the cell membrane. All three DNA origami configurations with 8 handles (“packed”, “patchy”, or “spread”) showed a strong decrease in the fraction of immobile tracks compared to the control condition that had 72 handles (Fig.2c,d). Notably, the 8-handle “packed” and “patchy” designs showed similar diffusion properties, while the “spread” design exhibited more immobile trajectories (Fig.2c-e). Interestingly, the “spread” design distributed handles uniformly along the same perimeter as the 72-handle configuration, resulting in identical bounded areas (38.5 nm × 36 nm) (Fig.1c, Fig.2b). Diffusion coefficient analysis of freely diffusive trajectories revealed that the “packed” design diffused fastest, followed by “patchy,” then “spread” and 72-handle designs (Fig.2d,f). This order corresponds directly to the PM area bounded by the DNA origami handles: “packed” designs covered the smallest area, followed by “patchy,” with “spread” and 72-handle designs covering equivalent areas. These results demonstrate that the specific spatial arrangement of lipid anchors, rather than simply the number of lipid anchors, determines pegboard-PM interactions. Since individual lipid anchors diffuse freely in the membrane, reduced pegboard diffusion reflects the PM’s inherent spatial organization, validating the usefulness of our approach for probing PM nanodomains.

**Figure 2.**
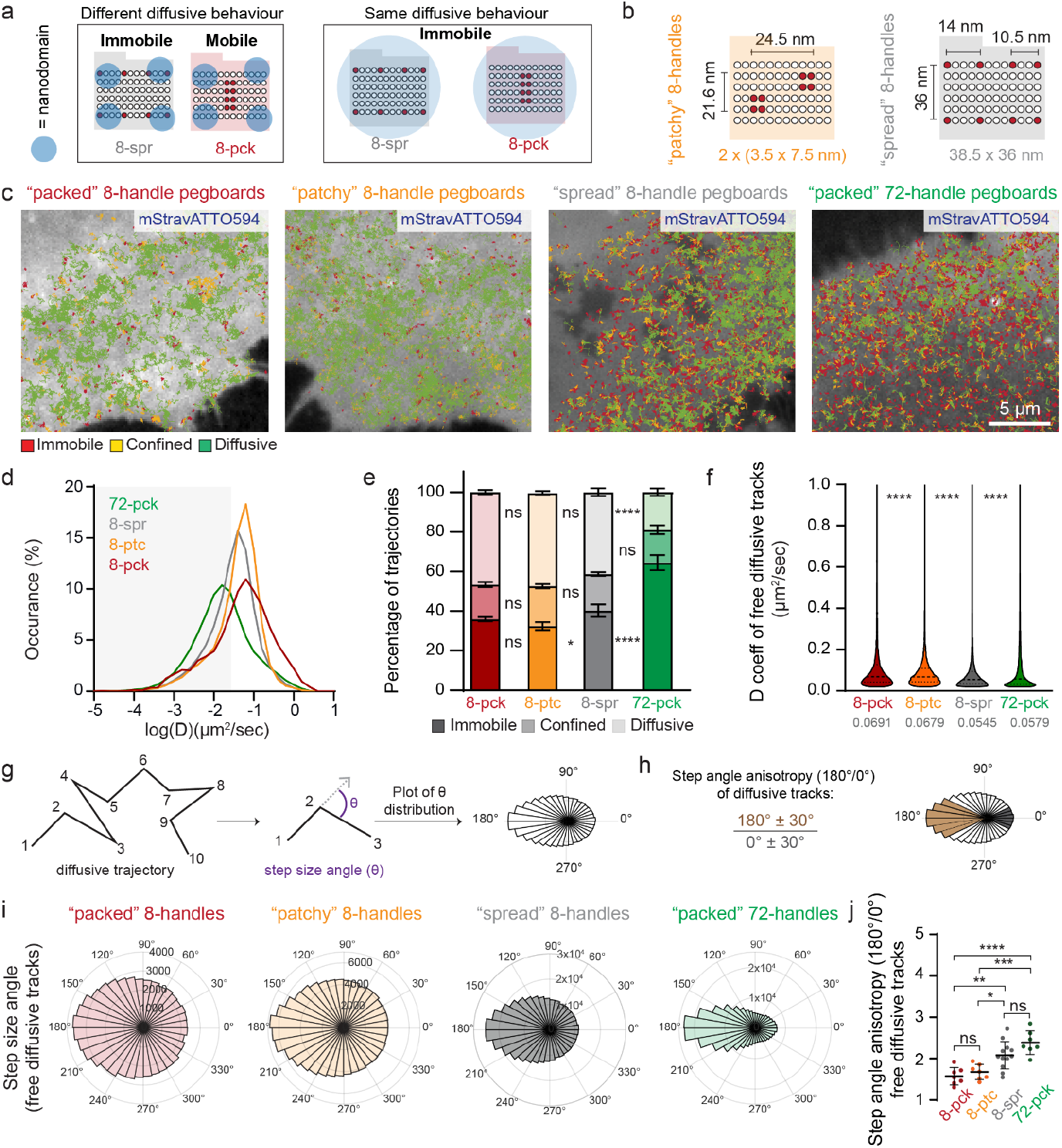
Spatial arrangement of lipid anchors influences DNA origami diffusion and anisotropy. **a**,**b)** Schematic of the tested hypothesis (a) and DNA origami designs (b). All origami displayed 8 handles for lipid anchors but varied in their handle arrangement. **c)** DNA origami trajectories overlaid on 6-FAM cell plasma membrane (gray). Trajectories are color coded to show their diffusion modes: diffusive (green), confined (yellow) and immobile (red). **c)** Distributions of the mean diffusion coefficient D computed from DNA origami trajectories. **d)** Fractions of tracked DNA origami undergoing free diffusion, confined diffusion or immobilization in the plasma membrane. **e)** Diffusion coefficient D for all free diffusive tracks of the DNA origami. **f**,**g)** Schematic of the step size angle and anisotropy analysis (adapted from^74^). **h)** Representative step size angle distribution for the diffusive tracks of the tested DNA origami. **i)** Step angle anisotropy of all the diffusive tracks. Data information: mean ± SEM (e,j), median and interquartile range (f). Statistics: n=5 (8-pck), n=7 (8-ptc), n=12 (8-spr), n=6 (72-pck), d) 2-way ANOVA with Tukey multiple comparisons test, e) number of diffusive tracks 13648 (8-pck), 26862 (8-ptc), 75901 (8-spr), 23228 (72-pck), Kruskal-Wallis test (non-parametric one-way ANOVA) with Dunn’s multiple comparisons test.. i) ordinary one-way ANOVA Tukey multiple comparisons test, each dot corresponds to the step angle anisotropy per single cell. Abbreviations: pck, packed; ptc, patchy; spr, spread.

### Spatial arrangement of lipid anchors affects the anisotropy of DNA origami diffusion

To further confirm that DNA origami pegboard diffusive properties reflect PM nanodomain interactions, we analyzed trajectory isotropy for free-diffusing and confined trajectories. This analysis captures transient confinement events that may be missed in global diffusion classification, where trajectories that appear to be overall free-diffusive could contain brief confined periods. To assess trajectory isotropy, we measured step-size angles for every diffusive trajectory and plotted distributions for each pegboard type (Fig.2g). We quantified anisotropy using “fold anisotropy”, meaning the ratio of steps with ∼180° reorientations to steps with ∼0° reorientations (straight travel) (Fig.2h). Higher anisotropy indicates greater confinement, as confined particles experience more frequent large reorientations and fewer forward steps, while more isotropic trajectories indicate free diffusion. The “packed” and “patchy” 8-handle pegboards exhibited broad step-size angle distributions and low fold anisotropy values (Fig.2i,j). Conversely, “spread” 8-handle and 72-handle pegboards showed narrower angle distributions and significantly higher fold anisotropy (Fig.2i,j). This increase in anisotropy correlates directly with the increased PM area bounded by the different designs. Our analyses of angles from confined DNA origami tracks revealed similar trends across all four designs (Fig.S3a,b), with significant differences between “packed” and “spread” 8-handle variants. These findings indicate that the PM contains a high density of nanodomains smaller than 20 nm, which are avoided by compact platforms but effectively trap larger-footprint designs covering ∼36 nm × 34 nm areas. The differential immobilization, diffusion coefficients, and anisotropy suggest that, while compact platforms primarily interact with individual nanodomains, larger platforms span multiple proximal nanodomains simultaneously, resulting in enhanced confinement (Fig.S3c).

### DNA origami tracks display diffusion-confinement cycles

To gain deeper insight into the organization of PM nanodomains, we extended our analyses beyond global diffusion behavior. Reasoning that DNA origami local confinements (i.e. immobilization and confined diffusion) reflect nanodomain organization, we focused specifically on characterizing these regions. To this end, we selected the 72-handle DNA origami design, which showed the greatest degree of confinement and the highest likelihood of engaging multiple nanodomains simultaneously. We performed sub-track analysis on long-duration trajectories (>50 frames) to resolve how confinement evolves in both space and time. This approach allowed us to identify local confined sub-regions within single trajectories, interspersed with free-diffusive periods, and to quantify changes in diffusivity along the path of individual probes (Fig.3a,b, see Methods for details). From the MSD analysis of each sub-track, we first computed the radius of confinement (R_conf_), which provides an estimate of the spatial extent over which the probe remains locally constrained. The R_conf_ revealed pronounced local confinement, with a mean radius of 48.02 ± 0.58 nm and with 33.47% of sub-tracks exhibiting confinement radii below the system’s spatial resolution (σ = 33 nm, Fig.3c), and only 3.34 % having radii above 100 nm. Notably, the time spent in confined states significantly exceeded time spent in free-diffusing states (Fig.3d). Together, these findings support a model in which individual DNA origami transiently engage multiple discrete, high-density nanodomain areas, resulting in complex and dynamic confinement behavior on the plasma membrane.

**Figure 3.**
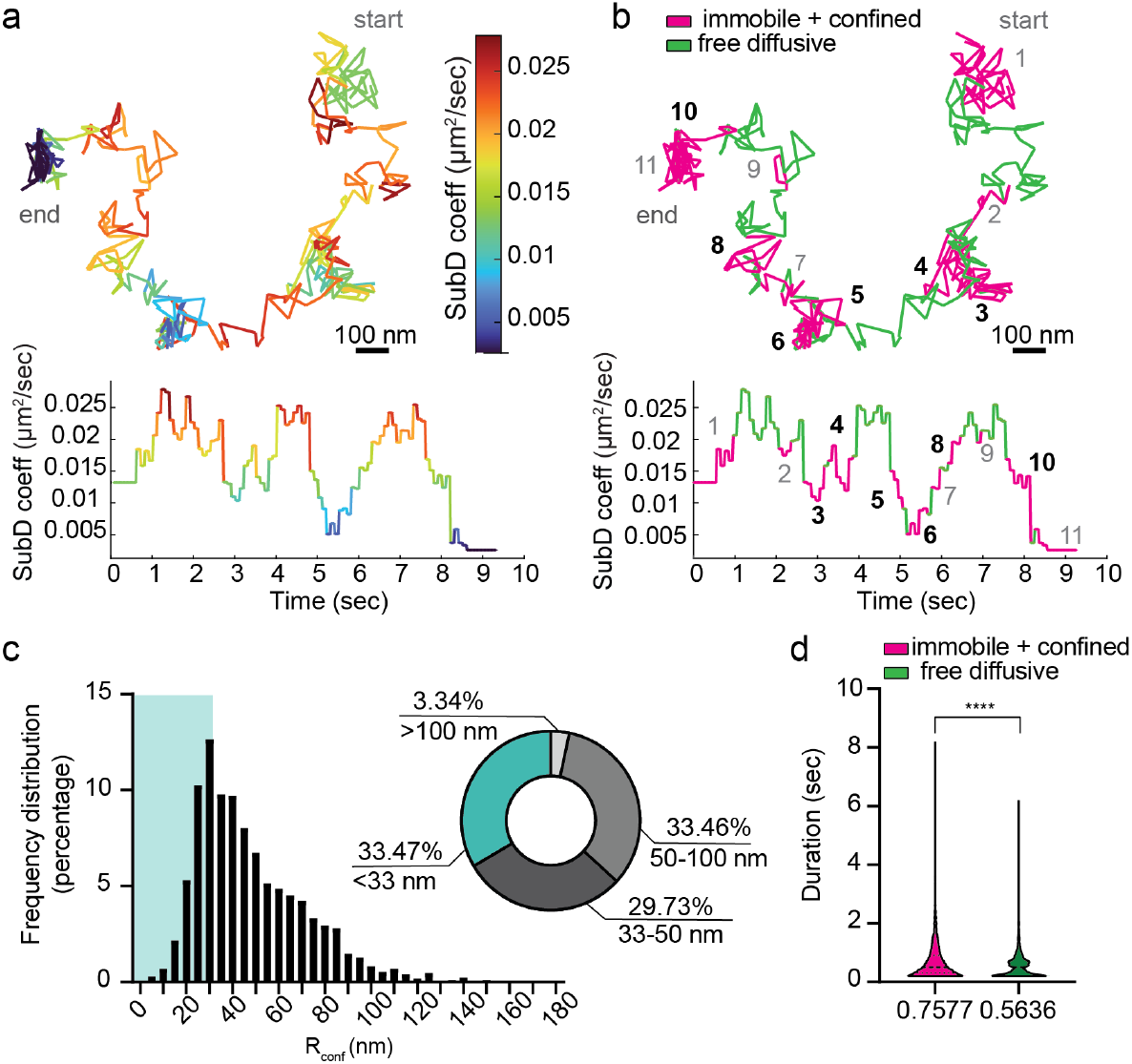
DNA origami tracks display diffusion-confinement cycles. **a)** Sub-track diffusion coefficient, D, as a function of time and space for a representative trajectory from a freely diffusing origami. **b)** Single trajectory and sub-track diffusion coefficient plot, color coded to show the sub-diffusion modes: free diffusion (green), confined diffusion and immobilization (magenta). Bolded numbers correspond to regions passing the selection criteria: the sub-track must be embedded between free diffusive sub-tracks and has a minimum length of 10 frames. **c**,**d)** Distribution of radii of confinement for origami experiencing confined diffusion or immobilization as a violin plot (c), pie plot and histogram as a percentage (d). Shadowed region in the plots highlights the values below our resolution limit (σ= 33 nm). **e)** Duration in seconds of the different sub-diffusion modes: free diffusion (green), confined diffusion and immobilization (magenta). Mean values are reported. Data information: median and interquartile range (d). Statistics: n=1450 (number of total trajectories longer than 50 frames), number of confined and immobile regions 4120 (magenta), and of free diffusing regions 2801 (green), n = 1709 (R_conf_ values). Unpaired Mann-Whitney t-test (d). Abbreviations: SubD, diffusion coefficient D computed on sub-tracks; R_conf_, radius of confinement.

### Membrane nanodomain stability partially rely on actin

PM nanodomains interact indirectly with the actin cytoskeleton and these interactions are crucial for forming and stabilizing nanodomains ^11,44,45^. Consequently, PM nanodomains are sensitive to the underlying cortical actin network ^8,12,46^ with actin filament destabilization by Latrunculin treatment inducing decreased sphingomyelin lipid entrapment ^46^ and reduced PM nanodomain size ^17^. Thus, we hypothesized that if our pegboards were interacting with PM nanodomains, actin cytoskeleton disruption should affect their diffusion. We performed live tracking of 72-handle pegboards on lipid anchor-decorated cells treated with and without LatrunculinA (LatA). Actin depolymerization mediated by LatA decreased the fraction of immobile 72-handle platforms by 23% (Fig.4a-c). Further analysis revealed that 72-handle pegboards on LatA-treated cells exhibited faster free diffusion than those on untreated cells (Fig.4d), while we observed no differences in fold anisotropy for free diffusive and confined tracks (Fig.S4a,b). These results demonstrate that the actin cytoskeleton significantly influences pegboard diffusion and support the hypothesis that DNA origami pegboards are trapped in PM nanodomains.

**Figure 4.**
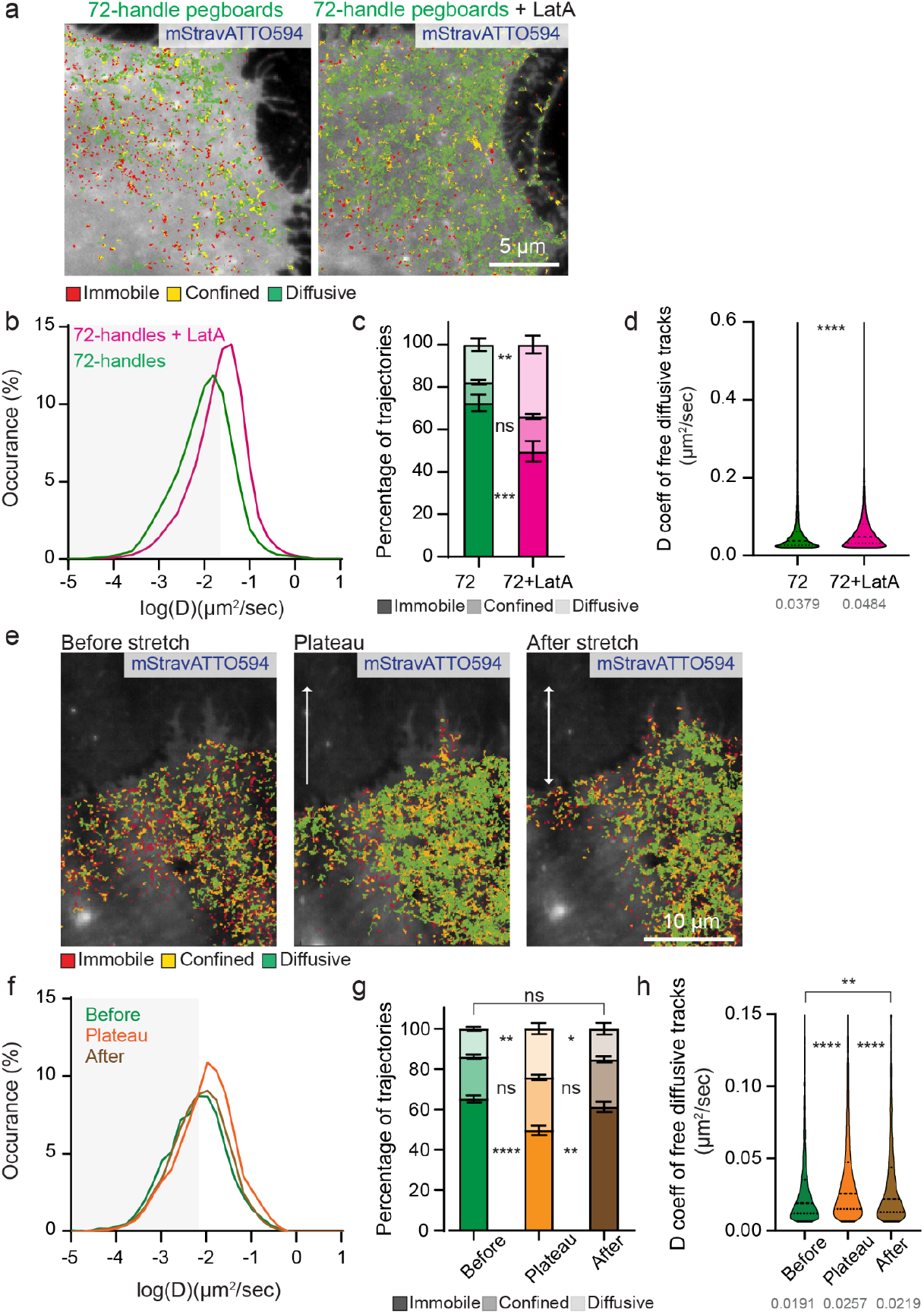
Membrane nanodomain organization is sensitive to actin and mechanical stimulation. **a)** Trajectories of 72-handle DNA origami diffusing on the cell membrane in the presence or absence of LatrunculinA (LatA) overlaid on 6-FAM cell plasma membrane (gray). Trajectories are color coded to show their diffusion modes: diffusive (green), confined (yellow) and immobile (red). **b)** Distributions of the diffusion coefficient D computed from the trajectories of 72-handle DNA origami with or without LatA. **c)** Fractions of tracked 72-handle DNA origami undergoing free diffusion, confined diffusion or immobilization in the plasma membrane with or without LatA actin destabilization. **d)** Diffusion coefficient D for all free diffusive tracks of 72-handles DNA origami in LatA presence or absence. **e)** Representative images of the 6-FAM positive cells superimposed with 72-handles DNA origami tracks before stretch, at the plateau and after stretch. Arrows indicate the stretching and relaxation direction. **f)** Distributions of the diffusion coefficient D computed from the trajectories of 72-handles DNA origami in all the conditions. **g)** Fractions of tracked 72-handles DNA origami undergoing free diffusion, confined diffusion or immobilization in the plasma membrane before stretch, at the plateau and after stretch. **h)** Diffusion coefficient D for all free diffusive tracks of the tested conditions. Data information: mean ± SEM (c,g), median and interquartile range (d,h). Statistics: n=6 (72-handles), n=8 (72-handles + LatA), n=6 (Before), n=4 (Plateau), n=4 (After); c,g) 2-way-ANOVA with Sidak multiple comparisons test, d) number of diffusive tracks 3789 (72-handles), 54126 (72-handles + LatA), Mann-Whitney test, h) number of diffusive tracks 4272 (Before), 6870 (Plateau), 2233 (After), 1-way-ANOVA with Holm-Sidak multiple comparisons test. Abbreviations: LatA, LatrunculinA actin destabilizer.

### Mechano-stimulation transiently regulates membrane nanodomain organization

Mechanical forces exerted on cells regulate membrane tension ^47^, impact the underlying actin cytoskeleton ^40,44^, and control membrane nanotopography by flattening caveolae or reshaping nanoscale membrane invaginations and evaginations ^48,49^. Moreover, actomyosin-generated forces influence PM nanodomains, controlling their formation and stabilization ^11,18,44^. To test how mechanical force regulates PM nanodomains, we combined cell stretching with DNA origami tracking using a stretching device we recently developed that is compatible with super-resolution microscopy and single-molecule tracking ^40^. We tracked 72-handle pegboards on lipid-decorated MEF cells before, during, and after 10% uniaxial stretch in 30 seconds. Stretching significantly increased the fraction of freely diffusive trajectories with concomitant decreased immobilization at peak stretch compared to unstretched conditions. Importantly, upon relaxation, the diffusive pegboard fraction was restored to pre-stretch levels (Fig.4e-g). Analysis of diffusion coefficients revealed higher speeds during stretching than before and after relaxation (Fig.4h), while no differences were observed in fold anisotropy for free diffusive and confined tracks (Fig.S4c,d). These results suggest that mechanical force can control PM nanodomain organization by decreasing nanodomain number or density. The rapid and reversible effects indicate an acute membrane or cytoskeletal mechanism rather than the effects due to slower processes such as cholesterol depletion ^2,50^.

## Discussion

Our studies show that immobilization and diffusion of our DNA origami pegboards on the PM is influenced by several factors: number and spatial arrangement of lipids on the DNA origami, actin destabilization, and mechanical forces. These findings demonstrate that DNA origami pegboards are an effective method for probing the nanoscale dynamic organization of the PM in living cells. SPT in concert with DNA origami has been previously used to study enzyme dynamics ^51^, membrane proteins ^52^ and to mimic biological membrane components ^53^. In addition, DNA origami have been patterned via lipophilic anchors to act on both model ^32–36^ and cell membranes ^37,38^. We further demonstrate the power of DNA origami to decipher PM nanoscale organization, the first use of DNA origami for this purpose. Our approach revealed that dynamic PM nano-organization is characterized by the presence of nanodomains of varying sizes, likely with a high density of small nanodomains (5-20 nm). Furthermore, we demonstrated that PM nanodomain organization depends only partially on the actin cytoskeleton and is reversibly altered by cellular stretching. These results establish DNA origami as a powerful tool for studying PM nanodomains while providing new insights into their density, size distribution, and dynamic organization.

We probed nanodomain density and size using various DNA origami pegboard designs. The 72-handle design (36 nm × 34 nm) showed the highest level of immobilization, suggesting entrapment of the DNA origami within either large (>36 nm) nanodomains or in multiple smaller, closely spaced ones. If all nanodomains were much larger than 36 nm, all the 8-handle DNA origami designs would behave identically, as all handles would interact with the same nanodomain, eliminating configuration-dependent effects on entrapment and diffusion. This is not what we observed. Indeed, the “packed” 8-handle pegboard (3.5 nm × 21.6 nm) was less immobilized than the “spread” 8-handle design. This indicates a high density of nanodomains smaller than 20 nm, through which the DNA origami “packed” design can avoid entrapment, whereas the “spread” 8-handle and 72-handle designs can effectively be trapped. Consistently, “packed” and “patchy” designs shared similar diffusion and anisotropy, whereas 8-handle “spread” pegboards behaved more like the 72-handle design (i.e., enhanced confinement, reduced diffusion, and higher immobilization) (Fig.S3c). Sub-track analysis data further support a model of high nanodomain density: more time is spent in confined or immobile states than in free diffusion, and 33% of confinement radii fall below the spatial resolution (<33 nm), with 97% under 100 nm (Fig.3). Our results align with previous reports of nanodomains ranging from 5–30 nm ^18^, or ∼20 nm ^19^, as inferred from homo-FRET, STED-FCS, and tracking of GPI-anchored proteins. Larger nanodomains in the 50–100 nm range have also been observed by direct super-resolution imaging of GPI-anchored proteins using STORM ^16^, NSOM ^10^, and PALM ^54^ techniques. However, unlike these approaches that target specific components, our method probes the entire plasma membrane beneath the DNA origami pegboard, providing new insights into the density and distribution of small (<20 nm) nanodomains, which appear to be both abundant and closely spaced.

This range in DNA origami diffusive behavior suggests that observed immobilizations and confinements reflect nanodomain presence rather than extracellular membrane component interactions ^8,12,14^. An alternative or complementary hypothesis is that membrane nanotopographies also influence DNA origami pegboard diffusion. Given the size and geometry of our DNA origami constructs, it is unlikely that they themselves generate significant membrane curvature or introduce nanotopographical features, as shown for single origami of similar dimensions on lipid model membrane ^33^. Rather, they are more plausibly acting as sensors of pre-existing membrane nanotopography. Consistent with this idea, mathematical simulations of protein diffusion in membranes with varying curvatures (from planar to vesicles of different radii) reveal that higher membrane curvature decreases the diffusion of embedded membrane proteins ^55^. In addition, molecular-dynamics simulations demonstrate that membrane proteins uniquely modulate their local lipid environment, enriching or depleting specific lipid components and creating thickness and curvature gradients ^43^. Beyond nanotopographies, the PM is compartmentalized into 50-200 nm domains created by interactions between the underlying actin cytoskeleton (fence) and transmembrane proteins (pickets) ^23,56^. This fence-and-picket organization causes anomalous diffusion of phospholipids and transmembrane proteins over long spatiotemporal scales, whereas free diffusion occurs within individual compartments and hop diffusion enables transitions between adjacent corrals ^23,57^. The observed differences between 8-handle “packed” and “patchy” versus 8-handle “spread” and 72-handle designs may reflect varying probabilities of hopping between actin-defined compartments. Larger or more extensively anchored DNA origami structures likely experience reduced intercompartmental mobility, particularly if corral sizes approach the membrane footprint of the pegboards. We observed changes in rates of DNA origami diffusion and immobilization following both actin destabilization and cell stretching. These findings support the hypothesis that DNA origami diffusive behavior could be regulated by incorporation into lipid nanodomains, interactions with PM nanotopography (e.g., membrane curvature), or the corralling effect, as all these processes are influenced by the actin cytoskeleton and mechanical stimuli ^11,44,48,49,58^.

We triggered actin-network disassembly in MEF cells by exposing them to acutely latrunculin A at a concentration that caused complete lamellipodium disassembly ^41^. LatA treatment reduced, but did not eliminate, DNA origami-lipid immobilizations and confinements on the PM, indicating that not all PM nanodomains were disassembled. These results suggest the coexistence of actin-dependent and actin-independent PM nanodomains. Actin-dependent nanodomains could form through actin filament connections to the PM ^8,11–14,44^, for example, through the ezrin/radixin/moesin proteins ^13,44^. Actin-independent nanodomains could be driven by PM nanotopographies ^43^, specific lipid compositions ^2^, or by the presence of transmembrane proteins not anchored to the cytoskeleton ^22^. Our DNA origami-lipid strategy provides high spatial resolution mapping without tracking individual nanodomain components (lipids or proteins), capturing all types of PM nanodomains in an unbiased manner, including those independent of the actin cytoskeleton. Our data suggest that approximately 40% of DNA origami-lipid immobilizations are actin-independent.

We previously developed a super-resolution-compatible stretching device to study mechanosensing in cells at the molecular level ^28,46^, showing that proteins such as talin undergo transient unfolding in integrin adhesions due to acto-myosin remodeling ^40^. Here, we leveraged this technology to investigate how mechanical forces affect PM nanodomain organization, also known to depend on actomyosin dynamics ^59^. Upon stretching, we observed a 10% decrease in immobile DNA origami tracks and a significant increase in free diffusion. This could reflect transient nanodomain disassembly due to disconnection from the cytoskeleton ^12,44,59^, increased membrane tension ^47,60,61^, altering PM nanotopography ^49,62^ or remodeling of cortical actin corralling effect ^58^. PM tension is a central regulator of mechanotransduction and mechano-protection ^60,63^, tightly linked to actin dynamics: increased tension inhibits actin polymerization, while reduced tension promotes it ^61,64^. PM tension causes caveolae flattening ^48,49^ and membrane nanotopography reshaping ^62,65^, and both events may contribute to increased DNA origami mobility. Importantly, diffusion changes were reversible upon relaxation (Fig.4e-h), consistent with transient PM deformations reported in myoblasts^48^ and fibroblasts^62,65^. Mechano-induced lipid raft disruption^16^, caveolae disassembly ^66,67^, and PM nanoscale reshaping ^62^, release PM-associated proteins (e.g. EHD2, Cav1 or ERM proteins)^2,47,66,67^ or recruit PM-bound proteins (e.g. the IRSp53 BAR protein)^62^, potentially triggering mechanotransduction^67^. Our findings show that mechanical stretch induces reversible plasma membrane nanodomain reorganization; this may be a mechanism for regulating nanodomain signaling activity.

PM nanodomains play a variety of roles in key cellular functions, yet, on the mechanical and molecular level, the PM remains largely poorly understood. Our DNA origami strategy complements the existing methods for studying PM nanodomains, enabling the detection of diffraction-limited membrane nanodomains. Overall, our data support a PM landscape model characterized by the co-existence of nanodomains larger than 20 nm and a high density of tiny nanodomains (5-20 nm) (Fig.S5). These nanodomains are partially stabilized by the underlying actin cytoskeleton, and mechanical forces can transiently remodel them. Our work provides a novel approach to accessing previously inaccessible information about these plasma membrane nanodomains, thus laying important groundwork for improving understanding of these essential dynamic nanostructures.

## Supporting information

Supplementary Table Sequences

## Acknowledgments

This work was supported by the National Institutes of Health (R21AI178200, R35GM125027, 1F32GM147967 to K.S.) and the National Science Foundation (DBI-1548297). We acknowledge financial support from the French Ministry of Research and CNRS, the Fondation pour la Recherche Médicale EQU202303016303 (to G.G.) ; the French National Research Agency: ANR-21-CE11-0004-01, CoCyNet (to G.G. and E.C.), ANR-22-CE13-0041, CryoNanoLam (to G.G.), the French government in the framework of the University of Bordeaux’s IdEx “Investiment for the Future” program/GPR BRAIN_2030, GPR LIGHT, the France BioImaging national infrastructure ANR-10-INSB-04. The authors thank Dr. Christophe Lamaze and Dr. Cedric Blouin for their critical reading of the manuscript. The authors also thank Leslie Biennen of C3 Science Communication for editing and feedback on the manuscript.

## Materials and methods

### Cell culture

Immortalized Mouse Embryonic Fibroblasts (MEF) were cultured in DMEM (Gibco) with 10% v/v Fetal Bovine Serum (FBS). On the day of the experiments, the cells were detached from the extracellular matrix using Trypsin/EDTA 0.05% for 2 min (Gibco). The trypsin was inactivated using soybean trypsin inhibitor (STI, Sigma) or with 10% (v/v) FBS DMEM. When trypsin was inactivated with 10% (v/v) FBS DMEM, cells were washed twice by consecutive cycles of centrifugation, supernatant removal, and resuspension in Ringer serum-free medium (150 mM NaCl, 5 mM KCl, 2 mM CaCl_2_, 2 mM MgCl_2_, 10 mM HEPES, 11 mM Glucose, pH 7.4) to avoid any trace of serum. 50k cells were seeded on human fibronectin (10 µg/ml, Sigma) coated 18 mm 1.5 H coverslips and kept in Ringer medium at 37°. Experiments were performed 3-5 hours after cell plating.

When treated with Latrunculin A (LatA) prior to live imaging cells were incubated for 5-10 minute with 1 µM LatA (Tocris).

### DNA origami

DNA origami pegboards were fabricated as described in previous works^28,68^. Briefly, DNA origami pegboards were designed using Cadnano (Fig.S6) ^69^. The pegboards were assembled from p8064 DNA origami scaffold (Tilibit), and chemically synthesized staple DNA oligonucleotides (Integrated DNA Technologies/IDT). Staples and scaffold were mixed at a molar ratio of 10:1 and folded in 12 mM Mg DNA origami folding buffer (12 mM Mg, 5 mM Tris, 1 mM EDTA). To fold the origami, the following steps were performed on a thermocycler:

1. hold at 65°C for 15’
2. drop to 60°C and hold for 1 hr
3. decrease temperature by 1°C and hold for 1 hr (repeat this step 19 more times, with a final hold temperature of 40°C)
4. hold at 25°C.

Folded origami were purified by PEG fractionation, then validated using agarose gel electrophoresis and negative stain transmission electron microscopy (TEM). All sequences used in this work can be found in the supplement (Table S1).

### Lipid anchor

Oligonucleotide synthesis was performed on an H8 automated synthesizer (K&A Labs, Germany) using the conventional phosphoramidite methodology on a one µmole scale. Trichloroacetic acid (TCA, Glen Research, USA) (3% in dichloromethane) was used for detritylation, 0.25 M 5-Benzylthio-1H-tetrazole (BTT, Glen Research) in dry acetonitrile was used as an activator, and oxidation was achieved using 0.02 M iodine in tetrahydrofuran/water/pyridine (Glen Research). The capping was achieved using a mixture of an acetic anhydride solution in THF (Cap A, Glen Research) and 10% 1-methylimidazole in THF/pyridine (Cap B, Glen Research). Pre-packed nucleoside 1000Å CPG (LINK, Scotland) and fast deprotecting β-cyanoethyl phosphoramidite monomers (Bz-dA, Ac-dC, dmf-dG, dT, Glen Research) were used to synthesize DNA. The monomers were dissolved in anhydrous MeCN (0.067 M) immediately prior to use. The fluorescein (6-FAM) was directly bound to the CPG solid support (3’-Fluorescein CPG), and the hexaethylene glycol (HEG) spacer was incorporated to isolate the origami docking oligonucleotide sequence from the lipidic segment of the lipid-DNA conjugate. All non-lipidic phosphoramidites and 6-FAM solid supports were purchased from (LINK).

Lipidic modifications of the oligonucleotide sequences were incorporated during the last cycle of the automated DNA synthesis with 2 previously reported custom-made phosphoramidites: one with a simple C18 saturated alkyl modification and the second one with a ketal modified uridine derivative ^70^.

The lipid-oligonucleotide sequences synthesized were: 5’ (C18)-TTTCTTCATTA–HEG-TTCCTCTACCACCTACATCACTT-(6 FAM)-3’; 5’ (ketal)-TTTCTTCATTA–HEG-TTCCT CTACCACCTACATCACTT-(6 FAM)-3’ used in combination with the 12-handles origami (Fig.S2), and 5’ (ketal)-TTTCTTCATTA–HEG-AAGATGAGGTAGATGGTT-(6 FAM)-3’ with all the other origami designs (Fig.1-4, Fig.S3-6). Cleavage and deprotection were performed according to supplier protocol. The crude oligonucleotide solution was concentrated under reduced pressure and redissolved in water (0.5 mL). The oligonucleotide sequences were finally dialyzed first against 50 mM NaCl and then twice against water. The oligonucleotide concentrations were determined from the absorbance value at 260 nm and the oligonucleotide’s epsilon (molar extinction coefficient). These values were calculated using the Integrated DNA Technology online oligo analyzer tool which uses the standard nearest neighbor method.

### Single lipid and DNA origami tracking

For single particle tracking with DNA-PAINT (DNA-PAINT-SPT), cells were incubated with lipid anchors (2 µM) diluted in Ringer medium (for single lipids) or in 10 mM MgCl2 Ringer medium (for DNA origami) at room temperature for 15 minutes and washed 3 times with the same dilution medium to remove free lipids not inserted in the plasma membrane. For single lipid tracking experiments (Fig.1, Fig.S1) 0.5-1 nM cy3b-Imager P3 (Massive Photonics) was added to the cells and acquired at room temperature without further incubation. For single-particle tracking of DNA origami (Fig.1-4), 2 nM biotinylated DNA origami was mixed with 1 µM mStravATTO594 for imaging. The mix, diluted in 10 mM MgCl2 Ringer medium, was added to the cell with lipid anchors. DNA origami and mStravATTO594 were used at a final concentration during acquisition of 1 nM and 0.5 µM, respectively. Cells were imaged at room temperature in an open chamber (Ludin chamber, Life Imaging Services, Switzerland) mounted on an inverted motorized microscope Nikon Ti equipped with a 100x 1.45 NA PL-APO objective, an automatic perfect focus system, and a Total Internal Reflection Fluorescence (TIRF) illumination module. The angle of the incident laser beam was adjusted to selectively illuminate the surface of the plasma membrane at the periphery of the cell, where the sample thickness allows a proper TIRF illumination. Cy3b imager P3 (for single lipid tracking) or mStravATTO594 (for DNA origami) were excited at 561 nm (5 mW at the objective), keeping the single molecule regime during multiple frames. The fluorescence was collected by the combination of a dichroic mirror and emission filters (D101-R561 and F39-61,7 respectively, Chroma, USA) and a sensitive EMCCD (Evolve, Photometric, USA). 5 streams of 4000 frames each were acquired. Fluorescein (6-FAM) (on the lipid anchor) was excited using a 488 nm laser. The acquisition was driven by Metamorph software (Molecular Device, USA) in streaming mode at 50 Hz, 20 ms exposure time (for the single lipids and DNA origami, Fig.S1d-f, Fig.1-4, Figs.S3 and S4) or 500 Hz, 2 ms exposure time (single lipids control experiment, Fig.S1h-j).

### DNA origami tracking on stretching devices

Stretching devices were built as previously ^40,71^. MEF cells culture and DNA origami labelling was done as for coverglass. The strain of the uniaxial stretch was calibrated using fluorescence beads (Tetraspeck, ThermoFisher) seeded on the PDMS substrate. As before, cells were imaged at room temperature on an inverted motorized microscope Nikon Ti equipped with a 100x 1.45 NA PL-APO objective, an automatic perfect focus system, and TIRF illumination module. mStravATTO594 were excited at 561 nm (12 mW at the objective) keeping the single molecule regime during multiple frames. Fluorescence data were collected using a combination of a dichroic mirror and emission filters (mEOS filter, Semrock FF02-617/73-25) and a sensitive a sensitive scientific complementary metal-oxide semiconductor sCMOS (ORCA-Flash4.0, Hammamatsu). Acquisition was driven by Metamorph software (Molecular Device, USA) in streaming mode. 2 streams of 4000 frames were acquired with a 50 ms exposure time for each step of the stretching protocol: 2 streams before, 2 streams at the stretching plateau and 2 streams at the relaxation. The uniaxial mechanical stretch applied was a single 10% stretch performed in 30 sec.

### Tracking analysis

Single molecules were localized and tracked over time using PALMTracer in MetaMorph ^72^. For stretching experiments, beads were used to register the localizations using the registration option within PALM Tracer. Under the experimental settings described above, and the basal camera noise measured experimentally, the resolution of the whole system on coverglass (microscope equipped with a EMCCD (Evolve) camera, pixel size of 160 nm) was 79.6 nm and on the stretching devices (microscope equipped with a sCMOS (ORCA-Flash4.0, Hammamatsu) camera, pixel size of 130 nm) was 71.2 nm. The resolution was determined to be 2.355 ***σ***_xy_, where ***σ***_xy_ is the pointing accuracy of the Gaussian fitting estimated with Thunderstorm ^39^. The trajectories were analyzed specifically inside the cell, thus removing the contribution of immobile detections on the coverglass, by using as a reference the segmented GFP image (derived from the alexa-488 of the lipid anchors). We analyzed trajectories lasting at least 10 points (i.e. 200 ms, for all experiments; 500 ms, for the stretching experiment) with a custom Matlab code, as previously done ^39,73^. The Matlab algorithm computes the mean square displacement (MSD) of the particles and the corresponding diffusion coefficient defined as the slope of the affine regression line of the MSD fitted for the four first values over time ^39^. We use step angle anisotropy to analyze the confinement of trajectories ^73,74^. First, step angles between consecutive steps of only diffusive trajectories and steps greater than the resolution limit are collected from each set of experiments. The distribution of the angles is drawn as a rose plot. The step angle anisotropy is then calculated as f_150-210_/60×360, where f_150-210_ is the fraction of angles between 150 and 210 degrees. For sub-trajectory analysis, we only analyzed tracks fulfilling these conditions: tracks longer than 50 frames, presenting local confinement (i.e. region of immobilization and confined diffusion) interspersed with free diffusive regions. For the main MSD analysis, a minimal length of 10 frames was required for each region of local confinement to be included in the analyses. Finally, the spatial extent of confinement was estimated by calculating the radius of confinement (R_conf_) as the square root of the MSD plateau value as previously described ^39,73^.

### Statistical analysis

All data were analyzed with Prism (GraphPad 7). Respective n values, data representation (i.e. median, mean, errors) and statistical test used are specified in the Figure legends. For all tests, the significance level was α = 0.05. Resulting p-values are indicated as follows n.s when p> 0.05; * when 0.01<P<0.05; ** when 0.001<P<0.01; *** when P<0.0001.

