## Supplementary Figures for "Using DNA origami to study nanoscale organization of plasma membranes"

**Supplementary Table**

Table S1 – Table reporting all sequences for DNA origami pegboards fabrication.

#### Supplementary Figures

### Figure S1

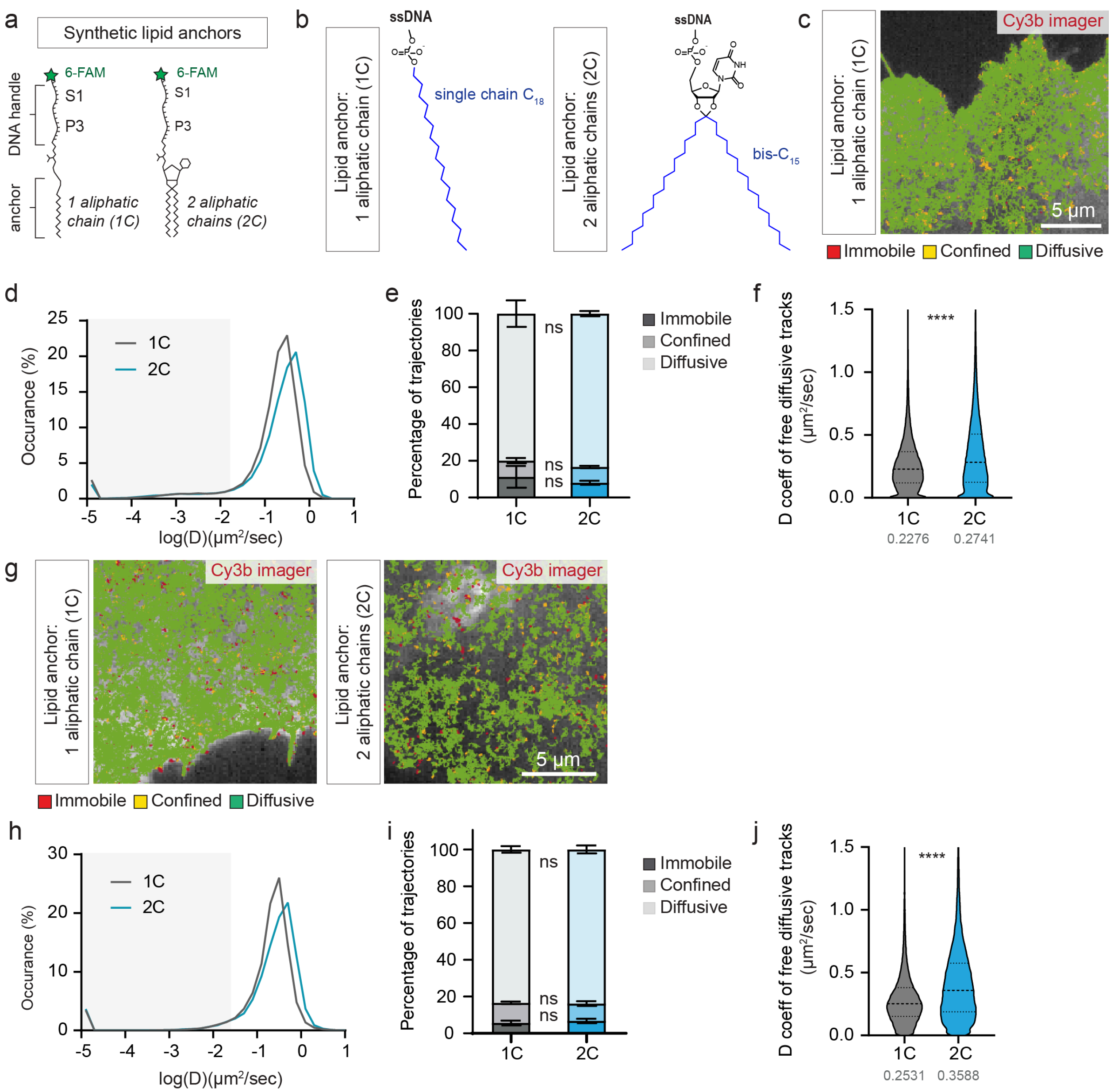


**Figure S1. Lipid anchors freely diffuse on the plasma membrane. a,b)** Graphical representation of lipid anchor structures (ssDNA sequence are reported in the method section). **c-f)** DNA-PAINT tracking of 1C and 2C lipid anchors acquired at 50Hz in MEFs. c) Trajectories of single lipid anchor 1C overlaid on 6-FAM cell plasma membrane (gray) – note that trajectories of single lipid anchor 2C are shown in Figure 1d. Trajectories are color coded to show their diffusion modes: diffusive (green), confined (yellow) and immobile (red). d) Distributions of the diffusion coefficient D computed from the trajectories of lipid anchor 1C and 2C. e) Fractions of tracked 1C and 2C lipids undergoing free diffusion, confined diffusion or immobilization in the plasma membrane. f) Diffusion coefficient D for all free diffusive tracks of 1C and 2C lipids. Data for 2C lipid anchors (d-f) are reported in Figure 1 for direct comparison in the presence/absence of DNA origami. **g-j)** DNA-PAINT-SPT of 1C and 2C lipids acquired at 500 Hz: g) Trajectories of single lipid 1C and 2C overlaid on 6-FAM cell plasma membrane (gray). Trajectories are color coded to show their diffusion modes: diffusive (green), confined (yellow) and immobile (red). h)

distributions of the diffusion coefficient D computed from the trajectories of lipid anchor 1C and 2C, i) fraction of tracked 1C and 2C lipids undergoing free diffusion, confined diffusion or immobilization in the cellular plasma membrane, j) diffusion coefficient D for all free diffusive tracks of 1C and 2C lipids.

Data information: mean ± SEM (e,i), median and interquartile range (e,i). Statistics: d-f) n=5, h-j) n=6, fe,i) 2-way-ANOVA with Sidak multiple comparisons test, f) number of diffusive tracks 120755 (1C), 132543 (2C), unpaired t-test, j) number of diffusive tracks 82764 (1C), 82519 (2C), unpaired t-test. Abbreviations: 1C, lipid anchor with one aliphatic chain; 2C, lipid anchor with two aliphatic chains; cy3b, cyanine 3b.

#### Figure S2

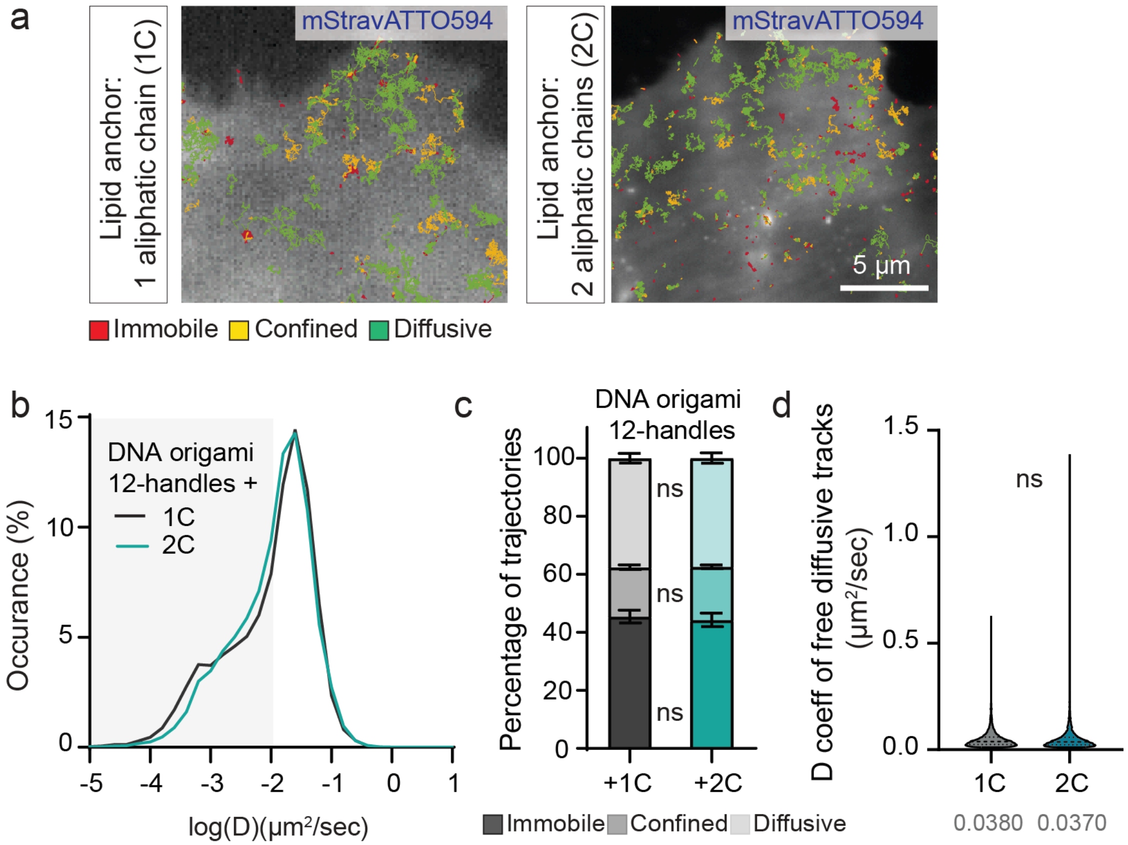


**Figure S2. DNA origami can be used as a probe for cell membranes. a)** Trajectories of 12-handles DNA origami, using either 1C or 2C lipid anchor, overlaid on 6-FAM cell plasma membrane (gray). Trajectories are color coded to show their diffusion modes: diffusive (green), confined (yellow) and immobile (red). **b)** Distributions of the diffusion coefficient D computed from the trajectories of 12-handles DNA origami using either 1C or 2C lipid anchor. **c)** Fractions of tracked 12-handles DNA origami combined with 1C or 2C lipids anchor undergoing free diffusion, confined diffusion or immobilization in the plasma membrane. **d)** Diffusion coefficient D for all free 12-handles DNA origami diffusive tracks combined with 1C or 2C lipids anchor.

Data information: mean ± SEM (c), median and interquartile range (d). Statistics: b-d) n=23 (1C) and n=18 (2C); c) 2-way-ANOVA with Sidak multiple comparisons test, d) number of diffusive tracks 13953 (1C), 13444 (2C), unpaired t-test. Abbreviations: 1C, lipid anchor with one aliphatic chain; 2C, lipid anchor with two aliphatic chains.

#### Figure S3

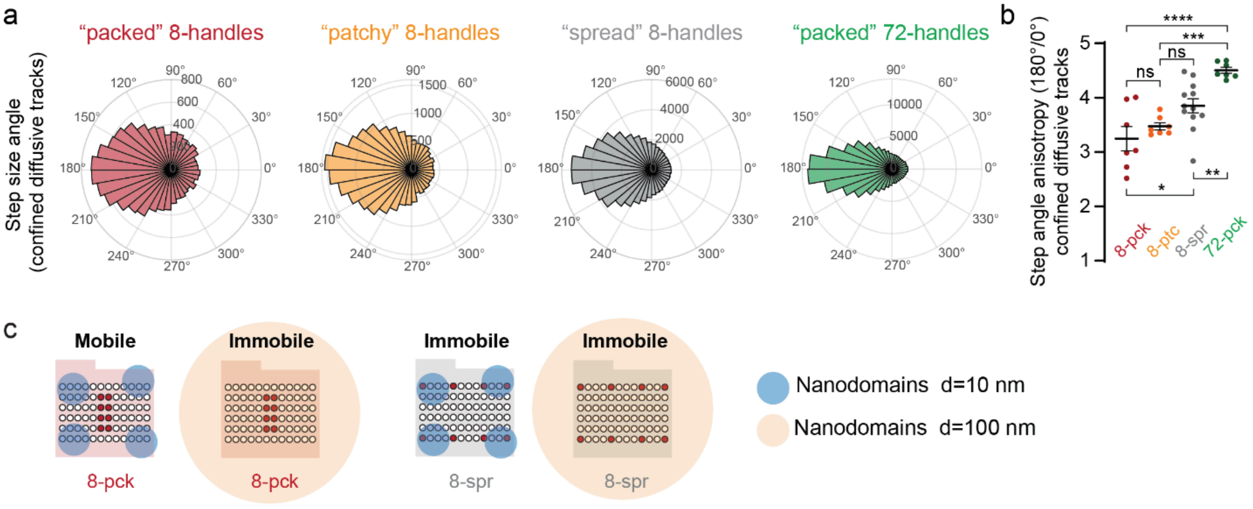


**Figure S3. The spatial arrangement of DNA origami handles for lipid anchors influences diffusion and anisotropy. a)** Representative step size angle distribution for the confined tracks of the tested DNA origami. **b)** Step angle anisotropy of all the confined tracks. **c)** Visual summary of the results.

Data information: mean ± SEM. Statistics: n=7 (8-pck), n=7 (8-ptc) n=12 (8-spr), n=7 (72-pck), b) ordinary one-way ANOVA Tukey multiple comparisons test, each dot corresponds to the step angle anisotropy per single cell. Abbreviations: pck, packed; ptc, patchy; spr, spread.

#### Figure S4

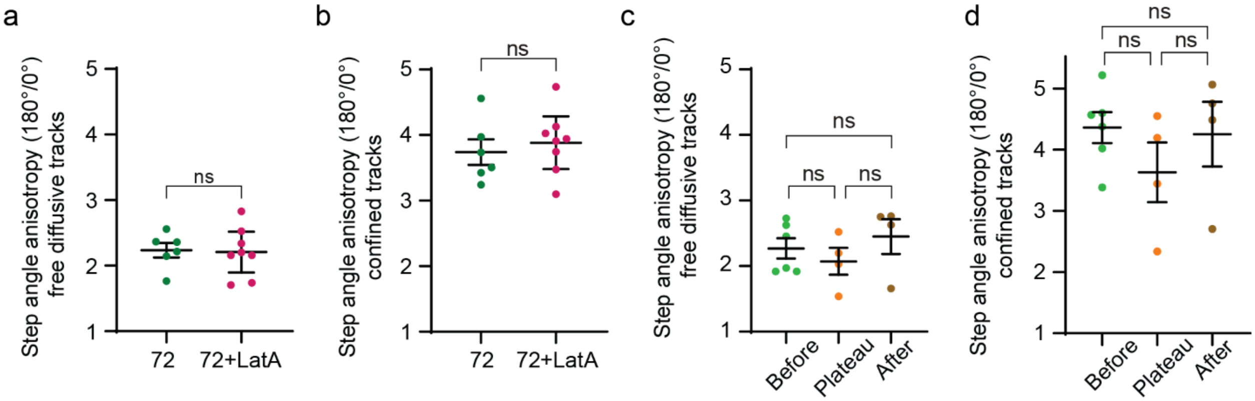


**Figure S4. Membrane nanodomains partially rely on actin and are sensitive to mechanical stimulation. a-d)** Step angle anisotropy of all the 72-handles DNA origami free diffusive tracks (a,c) and confined tracks (b,d) in the presence (+LatA) or absence of LatrunculinA and (a,b) before stretch, at the plateau and after stretch (c,d).

Data information: mean ± SEM. Statistics: n=6 (no LatA), n=8 (+ LatA), n=6 (Before), n=4 (Plateau), n=4 (After); (a,b) unpaired t-test, (c,d) Kruskal-Wallis test Dunn's multiple comparisons test, each dot corresponds to the step angle anisotropy per single cell. Abbreviations: LatA, LatrunculinA actin destabilizer.

#### Figure S5

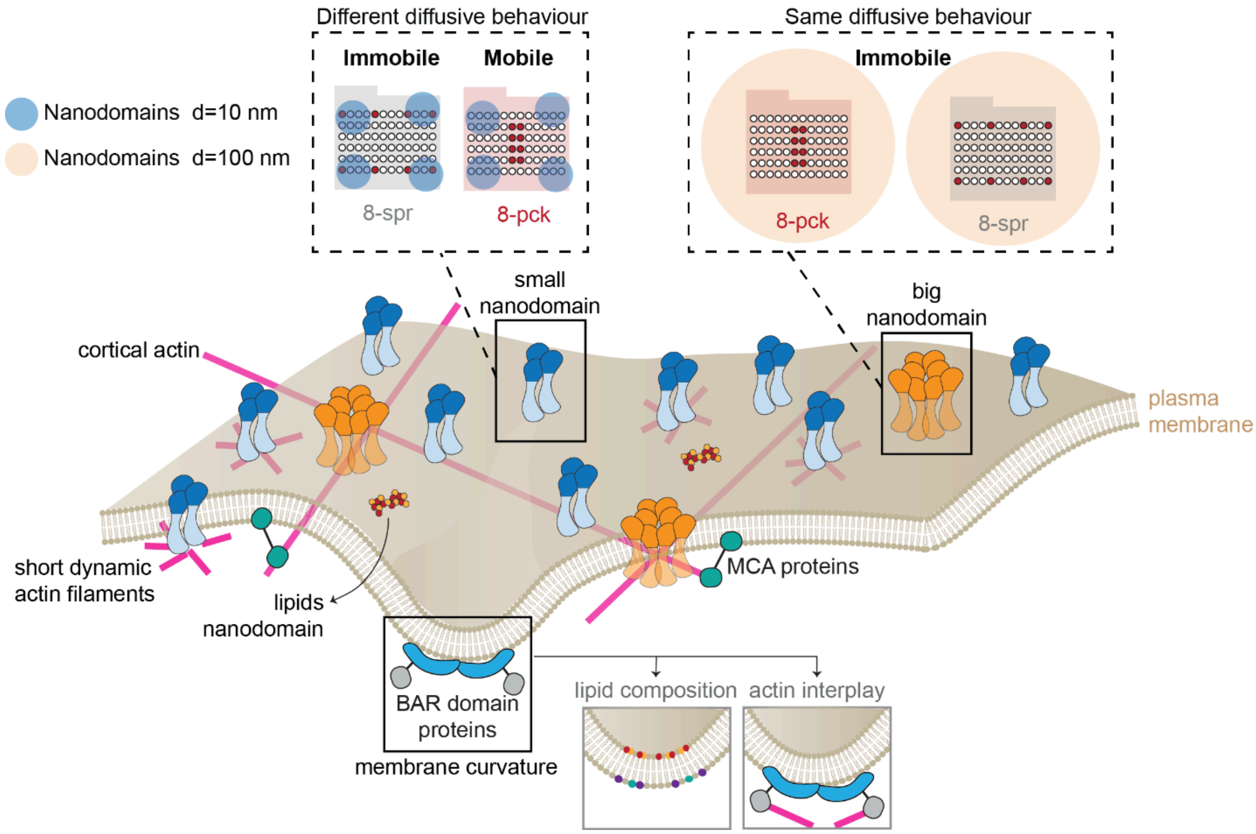


**Figure S5. DNA origami to study the nanoscale organization of the plasma membrane.** Schematic of the principle of DNA origami probing the plasma membrane nanodomains of different sizes (blue circles, nanodomains of 10 nm in diameter; light orange circles, nanodomains of 100 nm in diameter). DNA origami reveal a plasma membrane characterized by high density of small nanodomains (5-20 nm), dependent and independent on the actin cytoskeleton (magenta).

#### Figure S6

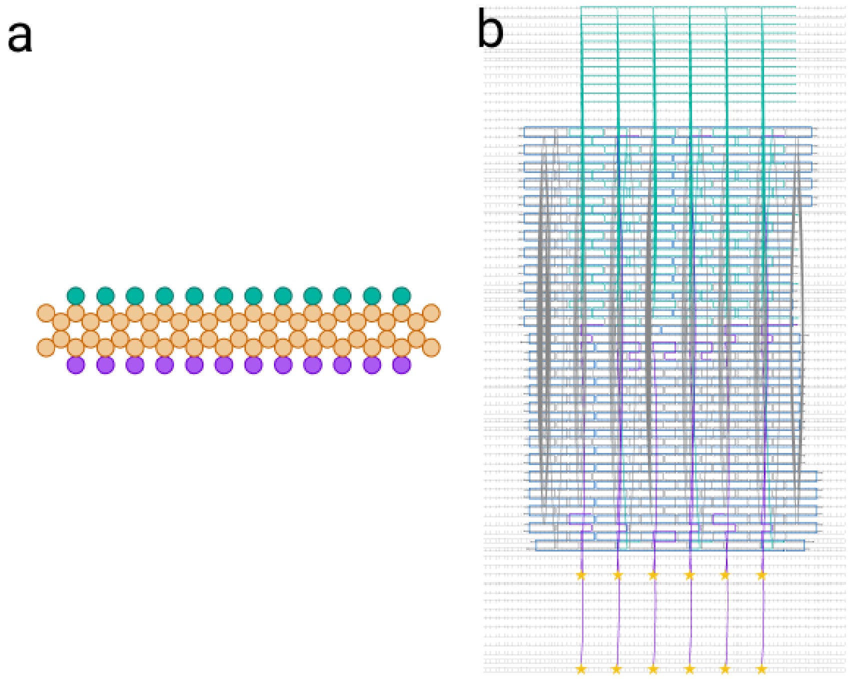


**Figure S6. Cadnano files depicting the cross sectional helix view (a) and the staple routing path (b).** Teal helices and staples represent staples bearing ssDNA handles for lipid anchors. Purple helices and staples represent biotinylated staples for conjugating mStrav-ATTO594. Biotin sites are denoted by stars and are located at the 5’ ends of the purple staples.
